# Early-life respiratory viral infection results in impairment of adult lung function

**DOI:** 10.1101/2021.09.20.461150

**Authors:** Laith H Harb, Patrick G Holt, Deborah Strickland, David Martino, Alexander N Larcombe, Anthony Bosco

**Affiliations:** Wal-yan Respiratory Centre, Telethon Kids Institute, Perth, Australia; Child Health Research Centre, The University of Queensland, Brisbane, QLD, Australia; University of Western Australia, Perth, Australia; Murdoch Children’s Research Institute, Department of Paediatrics, University of Melbourne, Melbourne, Australia; School of Population Health, Curtin University, Perth, Australia

**Author notes:** These authors contributed equally (Alexander N Larcombe, Anthony Bosco). Correspondence: Laith H Harb, Wal-yan Respiratory Centre, Telethon Kids Institute, Perth, Australia.

## Abstract

Respiratory viral infections in early-life are linked to the development of chronic lung diseases that persist into adulthood. The aim of this study was to develop a mouse model of early-life respiratory viral infection that would lead to impaired lung function in adulthood. BALB/c pups were infected at seven days of life with one of the following respiratory viruses: influenza A/Mem/1/71 “M71”, influenza A/Puerto Rico/8/34 “PR8” or attenuated mengovirus “Mengo”. Lung function and airways responsiveness (AHR) to methacholine were assessed seven weeks later, using the forced oscillation technique, and data were compared between male and female mice. PR8 infection was associated with significantly increased responsiveness to methacholine (for airway resistance, tissue damping, tissue elastance and hystersivity) for both sexes. M71 infection resulted in less severe responses especially in adult males. Early-life Mengo infection led to significantly higher responsiveness to MCh for males only (for airway resistance and tissue damping), suggesting sex dependant effects in lung function parameters measured. In summary, we have established a murine model where respiratory viral infection on day seven of life leads to AHR in adulthood. Importantly, the model recapitulates key variations in susceptibility related to sex and nature of viral pathogen that have previously been observed in human epidemiological studies. Our findings reveal new insights into the early origins of AHR and provide a tractable model system for future studies to unlock the mechanisms that determine pathogenesis.

## Introduction

Respiratory viral infections are the most common cause of illness among infants and are a major risk factor for the development of chronic lung diseases like asthma (1). Newborns are more susceptible to respiratory infections relative to older children and adults (2), with an average of five infections recorded for each child per year (3). The most common respiratory viruses encountered in early life are respiratory syncytial virus, rhinovirus, influenza virus and parainfluenza virus (1, 4). These infections, although typically self-limiting, can lead to chronic respiratory diseases in susceptible individuals. For example, infants who were hospitalised with RSV bronchiolitis in the first year of life displayed increased sensitivity to allergens and were more susceptible to developing subsequent asthma (5). Moreover, the association between early life respiratory infection and subsequent asthma may be even stronger for RV (6, 7) especially in combination with atopy (7, 8). Whilst the role of respiratory viral infections in asthma development is supported by a series of findings from longitudinal studies and experimental animal models, it is currently not known if these viruses damage the developing airways leading to disease or alternatively unmask individuals with a pre-existing susceptibility that were already on a trajectory towards disease (9).

Indeed, the first years of life are a crucial period of heightened plasticity where gene-environment interactions can program susceptibility to a range of chronic inflammatory diseases (10). Gollwitzer and Marsland suggested that an activation threshold must be overcome before a Th1 immune response can be effectively mounted (11). This threshold is affected by many exogenous factors. For example, mode of birth, use of antibiotics, mode of feeding (breast milk or formulae), stress, smoking, obesity, and malnutrition are known to impair development of the immune system during the prenatal and perinatal period, and in turn lead to ineffective host immune responses to pathogens (11). Conversely, exposure to microbes and their products, modify immune response patterns and modulate risk for the development of allergies and asthma (12). Exogenous factors can therefore on the one hand provide protection from disease by driving immune development or alternately on the other hand can be detrimental by promoting immune dysregulation and excessive inflammation.

The biological processes that control immune development operate in parallel to mechanisms that control lung development. The lung develops in five separate stages: embryonic, pseudoglandular, canalicular, saccular, and alveolar. While mice are born during the saccular stage and humans during the alveolar stage, murine models can be utilised to provide important insights into the role of infection/inflammation in lung development (13). Viral infections induce the production of a plethora of cytokines, chemokines and growth factors that recruit immune cells into the lung and damage local tissues (14).

Sex hormones influence the development of the immune and respiratory systems, which in turn leads to disparities in lung function and host responses to infections (15). Receptors for male and female sex hormones are expressed during foetal development, shaping lung anatomy and physiology (16). Androgen receptors are known to control branching morphogenesis, whilst estrogen receptors control lung maturation (17). Sex hormones also modulate the innate and adaptive arms of the immune system and determine the efficacy of pathogen clearance (18). For example, the female sex hormone estrogen enhances immune responses and promotes pathogen clearance (19). Moreover, neutrophils isolated from healthy women show improved survival and function, when compared to healthy male neutrophils (20). Alveolar macrophages are found in greater numbers and function more effectively in females, leading to more effective clearance of respiratory pathogens (21). However, androgens can induce the production of inflammatory cytokines (15), causing tissue injury and damage, resulting in increased risk of recurrent infections (22). Overall, sex hormones play important roles in the development and function of the immune and respiratory systems, which in turn may lead to long-term impacts of early life viral infections.

In this study, we established a murine model to determine if respiratory viral infections in early life leads to lung function deficits in adulthood. Influenza M71 and PR8 were chosen as representatives of mild and severe infections, respectively (23). Attenuated mengovirus was selected because it exhibits similar tropism to human rhinovirus and readily infects rodents (24). Experiments were performed in BALB/c mice because they display a Th2 bias and are susceptible to the development of chronic lung diseases (25). Responses were compared between male and female mice to examine the role of sex in disease development. Finally, a subset of the experiments was repeated in a second animal house facility to demonstrate that the experimental model was robust and reproducible.

## Methods

### Animals

Time-mated female BALB/c mice were purchased from the Animal Resources Centre (ARC) (Murdoch, WA, Australia) and delivered to the Telethon Kids Institute Bioresources Centre on day 15 of pregnancy. The animals were housed in individually ventilated cages (Sealsafe, Tecniplast, Buguggiate, Italy) on non-allergic, dust-free bedding (Shepherd Specialty Papers, Chicago, IL, USA). Animals were housed under a 12-hour day/night cycle. Food and water were provided *ad libitum*. Dams were left to give birth without any disruptions. All procedures were approved by the Telethon Kids Institute Animal Ethics Committee (Project #335).

### Viruses

Mengovirus is a serotype of encephalomyelitis virus and is part of the Cardiovirus species in the Picornaviridae family. Attenuated mengovirus (_V_MC_0_) was prepared as previously described (26). Influenza A/Mem/1/71 (H3N1) is a reassortment strain of the Influenza A family, comprising haemagglutinin of Influenza A/Mem/1/71 and neuraminidase of Influenza A/Bellamy/42. Influenza M71 was prepared as previously described (27). Influenza A/Puerto Rico/8/34 (H1N1) is a mouse-adapted Influenza strain. The virus was prepared as previously described (28).

### Infection

BALB/c litters were randomised into four experimental groups on day seven of life: influenza A/Mem/1/71 (M71), influenza A/Puerto Rico/8/34 (PR8), attenuated mengovirus (Mengo) or phosphate buffered saline as a control (Ctrl). On this day, pups were sexed, and administered virus at the following doses: 10^5^ plaque-forming-unit (pfu) of Mengovirus, 10^3.8^ pfu of Influenza A/Mem/1/71, 1.5 median-tissue-culture-infectious-dose (TCID_50_) of Influenza A/Puerto Rico/8/34. Inoculum was delivered in a volume of 10μL, intranasally via a pipette as previously described (29).

### Body weight measurements

Body weights were determined for all animals on day of infection and on the day lung function studies were performed. The delta weights were subsequently calculated for each experimental group and sex.

### Broncho-alveolar lavage (BAL) collection and processing

A random subset of male and female mice from each experimental group (2-11 per group) were sacrificed three-, five- and fourteen-days post-infection and the lung of each experimental subject was washed three times with 500μL of chilled phosphate-buffered saline (PBS) to obtain bronchoalveolar lavage fluid (BAL). BAL was centrifuged at 200 G for 4 minutes to obtain the cell pellet and supernatant. Resuspended cells were stained with trypan blue and total cell counts obtained using a haemocytometer.

### Lung function at functional residual capacity (FRC) and airway hyperresponsiveness (AHR)

At eight weeks of age, lung function was measured using the forced oscillation technique (flexiVent; Scireq, Montreal, Canada). An anaesthetic solution comprised of 0.4mL of ketamine (Troy Laboratories, Glendenning, NSW, Australia), 0.1mL xylazine (Troy Laboratories, Glendenning, NSW, Australia) and 0.5mL of saline was made and 1% of the total body weight was administered via intra-peritoneal route. Two-third of the anaesthetic solution was used to induce a surgical plane of anaesthesia, before an endotracheal tube (length = 1.0cm, internal diameter = 0.086cm) was inserted into the trachea and silk ligature was used to secure the tube. Once the animal was placed on the ventilator, the remaining third of the anaesthetic solution was given and the animals were then ventilated at 450 breaths/minute with a tidal volume of 8mL/kg and a positive end expiratory pressure of 2 cmH_2_O. A snapshot perturbation was performed once the animal was on the ventilator. A second snapshot perturbation was performed five minutes later. The single compartment model was fitted to the second perturbation from which resistance (R), elastance (E), and compliance (C) were extracted. Lung volume history was standardised via two deep inflations to a transrespiratory pressure of 20cm H_2_O, performed a minute apart (30). A third deep inflation was conducted a minute later, from which compliance (C), specific compliance (Cs), Vmax and the shape of the deflationary arm of the lung volume (K). Vmax is the highest volume at a transrespiratory pressure of 20 cmH_2_O. The compliance was calculated as the volume at 8 cmH2O minus the volume at 3 cmH2O divided by 5 (30). The specific compliance was calculated as the compliance divided by the volume at 3 cmH2O. The shape of the deflation arm of the PV loop (K) was calculated as the volume at 10 cmH2O divided by the volume at 20 cmH2O. Respiratory system impedance (Z_rs_) was then measured (one forced oscillation technique (FOT) measurement per minute, for five minutes) before the constant phase model was fit to Z_rs_. This allows for subsequent calculation of airway resistance (R_aw_), tissue damping (G) tissue elastance (H) and hystersivity (η). Hystersivity was calculated as G/H (31). Five measurements were made at functional residual capacity prior to a 10 second saline aerosol followed by five more measurements of Z_rs_. The animals were then subjected to increasing doses of methacholine (0.1, 0.3, 1, 3, 10, 30mg/mL) (a broncho-constrictor; acetyl-β methacholine chloride; Sigma-Aldrich) delivered by an ultrasonic nebuliser. Each methacholine aerosol lasted for 10 seconds. After each methacholine aerosol, five FOT measurements were collected and the maximum response for each measurement was used for subsequent analysis. This allowed for the construction of dose-response curves. Methacholine sensitivities were measured as the concentration of methacholine required to increase R_aw_, G, H and ɳ by 25% for each subject from saline. Then the values were averaged for each sex and experimental group and difference from control was calculated.

### Repeatability – Follow-up study

During the course of this study the Telethon Kids Institute Bioresources Centre which houses our animal holding facilities relocated to a new building. We endeavoured to determine if the relocation had any impact on outcomes. We focused on influenza M71 and influenza PR8 as representatives of mild and severe infection, respectively. We again measured lung function at functional residual capacity and responsiveness for MCh between the two locations.

### Statistics

Before performing ANOVA, we assessed our data sets for homoscedasticity and normality using Levene’s homogeneity of variance test and Shapiro-Wilk normality test, respectively. Unless otherwise stated, two-way ANOVA was used to analyse the data. A linear maximum likelihood model was fitted to the lung function data to allow comparison between experimental groups and determine statistical significance. Tukey multiple comparison test was used to identify significant differences between experimental groups and sexes. *P* < 0.05 was considered significant. For brevity, difference between baseline and the highest MCh dose will be considered significant for all lung function parameters. Statistical analyses were performed in GraphPad PRISM 7 (GraphPad Software, San Diego, California USA) or R (v4.0.4). Data are shown as mean ± SD.

## Results

### Variations in Body weight

To determine if the viral infection hindered somatic growth, we calculated the difference between body weights (i.e. delta weights) on the day of lung function study versus the day of infection. The data showed that the delta weights were not different between treatment groups (virus versus controls; p = 0.296), however delta weights were greater for males compared to females (p < 0.001) regardless of treatment. As noted above (see methods), we repeated a subset of the experiments in a follow-up study, and observed the same trend, whereby early-life viral infections did not affect delta body weights (p = 0.779). However, delta weights were again greater for males compared to females (p < 0.0001).

### Cellular Inflammation in BAL

At three days post infection with influenza M71, both male (p < 0.01) and female (p < 0.05) mice had significantly more total cells in their BAL compared with controls (Figure 1). There was no significant increase in total cells for either sex infected with Influenza PR8 or Mengo at this timepoint (Figure 1, p > 0.05 in all cases). There was no significant effect of any infection on total cells in BAL on days 5 or 14 post infection for either sex (Figure 1B and 1C, p > 0.05 in all cases). Differential cell counts revealed that macrophages were the most abundant cell type (>95%) across all experimental groups and sexes (data not shown).

**Figure 1:**
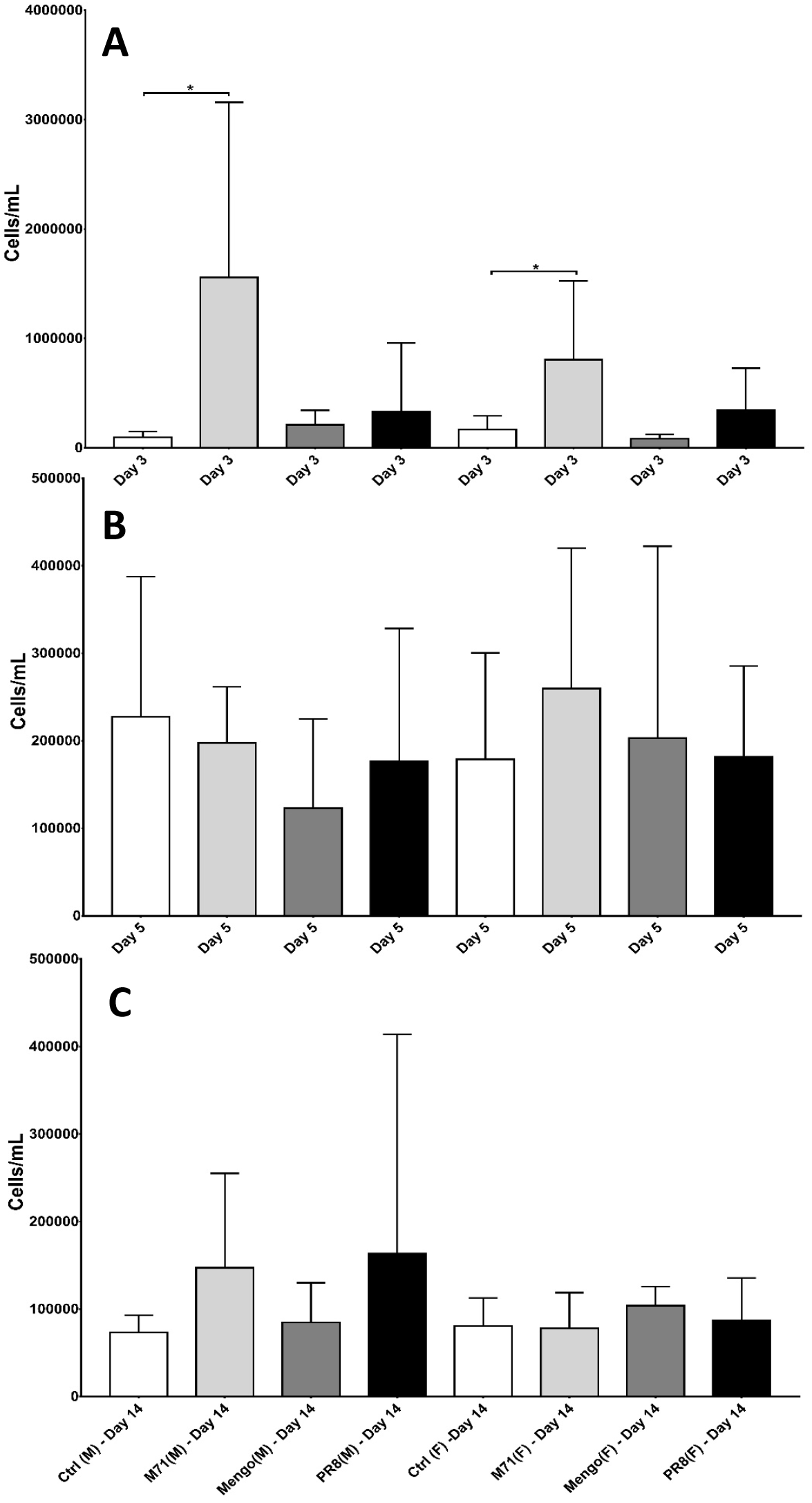
Total cell count for Ctrl (white bars), M71 (light grey bars), Mengo (dark grey bars) and PR8 (black bars) split by three (A), five (B) and fourteen (C) days post infection. Male mice are labelled by (M), and female mice are labelled by (F). The following numbers refers to the number of mice in each experimental group: Ctrl (M – Day 3: n = 6; M – Day 5: n = 11; M – Day 14: n = 10; F – Day 3: n = 8; F – Day 5: n = 7; F – Day: n = 6), M71 (M – Day 3: n = 3; M – Day 5: n = 5; M – Day 14: n = 9; F – Day 3: n = 3; F – Day 5: n = 3; F – Day 14: n = 7), Mengo (M – Day 3: n = 5; M – Day 5: n = 6; M – Day 14: n= 8; F – Day 3: n = 4; F – Day 5: n = 3; F – Day 14: n = 2) and PR8 (M – Day 3: n = 7; M – Day 5: n = 7; M – Day 14: n = 10; F – Day 3: n = 5; F – Day 5: n = 6; F – Day 14: n = 8). Data are mean ± SD. *N.B*. different scales for each bar graph.

### Resistance, elastance and compliance of the lung

We performed the snapshot perturbation technique to assess the lung as a single component providing data on resistance (R), elastance (E) and compliance (C) (32). Viral infections had no effect on R (p = 0.535), E (p = 0.685) or C (p = 0.644) for male or female mice (Table 2). Moreover, there was no effect of sex on these same parameters (R, p = 0.287, E p = 0.131, C, p = 0.097). There was no significant interaction between sex and early-life viral infection on any of these outcomes (p > 0.207 in all cases). We observed the same findings in the follow-up study, where viral infection had no effect on R (p = 0.306), E (p = 0.469) or C (p = 0.438) for male or female mice (Table 3), and there was also no significant interaction between sex and early-life viral infection on these parameters (R, p = 0.396, E, p = 0.312, C, p = 0.148) for male or female mice.

**Table 1:** Delta weights calculated as the difference in body weight between day of lung function study and first day of infection. Each data entry is classified as follows: mean ± SD (n). An asterisk (*) denotes a p < 0.05. *N.B*. Mengo was not replicated in the follow-up study.

| Delta weights – initial study |  |  |  |  |
| --- | --- | --- | --- | --- |
|  | Ctrl | M71 | PR8 | Mengo |
| **Male\*** ($p < 0.001$) | 16.54 ± 0.44 (11) | 17.16 ± 0.72 (23) | 16.46 ± 1.69 (23) | 16.62 ± 1.02 (19) |
| **Female\*** ($p < 0.001$) | 13.44 ± 1.3 (21) | 13.71 ± 0.57 (14) | 14.15 ± 1.43 (12) | 14.09 ± 0.3 (8) |
| Delta weights – follow-up study |  |  |  |  |
|  | Ctrl | M71 | PR8 | - |
| **Male\*** ($p < 0.001$) | 15.33 ± 0.48 (8) | 16.62 ± 0.2 (15) | 15.61 ± 0.72 (8) | - |
| **Female\*** ($p < 0.001$) | 13.02 ± 0.27 (6) | 13.79 ± 1.48 (6) | 13.63 ± 0.28 (7) | - |

**Table 2:** Snapshot perturbation parameters extracted for each sex and experimental group from the initial study. Data presented are mean ± SD.

|  | **Ctrl-F**<br>**(n = 10)** | **M71-F**<br>**(n = 7)** | **Mengo-F**<br>**(n = 5)** | **PR8-F**<br>**(n = 5)** | **Ctrl-M**<br>**(n = 4)** | **M71-M**<br>**(n = 13)** | **Mengo-M**<br>**(n = 10)** | **PR8-M**<br>**(n = 12)** |
| --- | --- | --- | --- | --- | --- | --- | --- | --- |
| Resistance (R)<br>cmH<sub>2</sub>O.s/mL | 1.40 ±<br>0.14 | 1.24 ±<br>0.18 | 1.28 ±<br>0.07 | 1.42 ±<br>0.22 | 1.28 ±<br>0.14 | 1.29 ±<br>0.25 | 1.27 ±<br>0.06 | 1.30 ±<br>0.25 |
| Elastance (E)<br>cmH<sub>2</sub>O/mL | 38.97 ±<br>5.93 | 33.89 ±<br>4.40 | 36.57 ±<br>1.71 | 38.80 ±<br>5.68 | 33.92 ±<br>2.92 | 35.80 ±<br>6.05 | 34.60 ±<br>2.18 | 35.54 ±<br>6.24 |
| Compliance (C)<br>mL/cmH<sub>2</sub>O | 0.03 ±<br>0.003 | 0.03 ±<br>0.005 | 0.03 ±<br>0.001 | 0.03 ±<br>0.003 | 0.03 ±<br>0.002 | 0.03 ±<br>0.004 | 0.03 ±<br>0.002 | 0.03 ±<br>0.004 |

**Table 3:** Snapshot perturbation parameters extracted for each sex and experimental group from the follow-up study. Data presented are mean ± SD.

|  | **Ctrl-F**<br>**(n = 6)** | **M71-F**<br>**(n = 6)** | **PR8-F**<br>**(n = 8)** | **Ctrl-M**<br>**(n = 7)** | **M71-M**<br>**(n = 14)** | **PR8-M**<br>**(n = 7)** |
| --- | --- | --- | --- | --- | --- | --- |
| Resistance (R)<br>cmH<sub>2</sub>O.s/mL | 1.27 ± 0.13 | 1.32 ± 0.18 | 1.39 ± 0.33 | 1.24 ± 0.09 | 1.11 ± 0.08 | 1.20 ± 0.19 |
| Elastance (E)<br>cmH<sub>2</sub>O/mL | 32.21 ± 4.26 | 34.14 ± 5.25 | 37.10 ± 10.58 | 31.38 ± 3.14 | 28.82 ± 2.82 | 30.03 ± 3.33 |
| Compliance (C)<br>mL/cmH<sub>2</sub>O | 0.03 ± 0.003 | 0.03 ± 0.003 | 0.03 ± 0.005 | 0.03 ± 0.003 | 0.03 ± 0.003 | 0.03 ± 0.003 |

### Pressure-volume (PV) curves and associated measurements

PV curves can be used to characterise functional changes in the lung (32, 33). The PV curves of infected male and female mice showed an upward shift when compared to control (Figure 2A and 2B). In the follow-up study, the PV curves of male M71 infected mice shifted upwards, while the PV curve of PR8 male mice were similar to control (Figure 2C). In female mice, the M71 and PR8 PV curves were also similar to control (Figure 2D).

**Figure 2:**
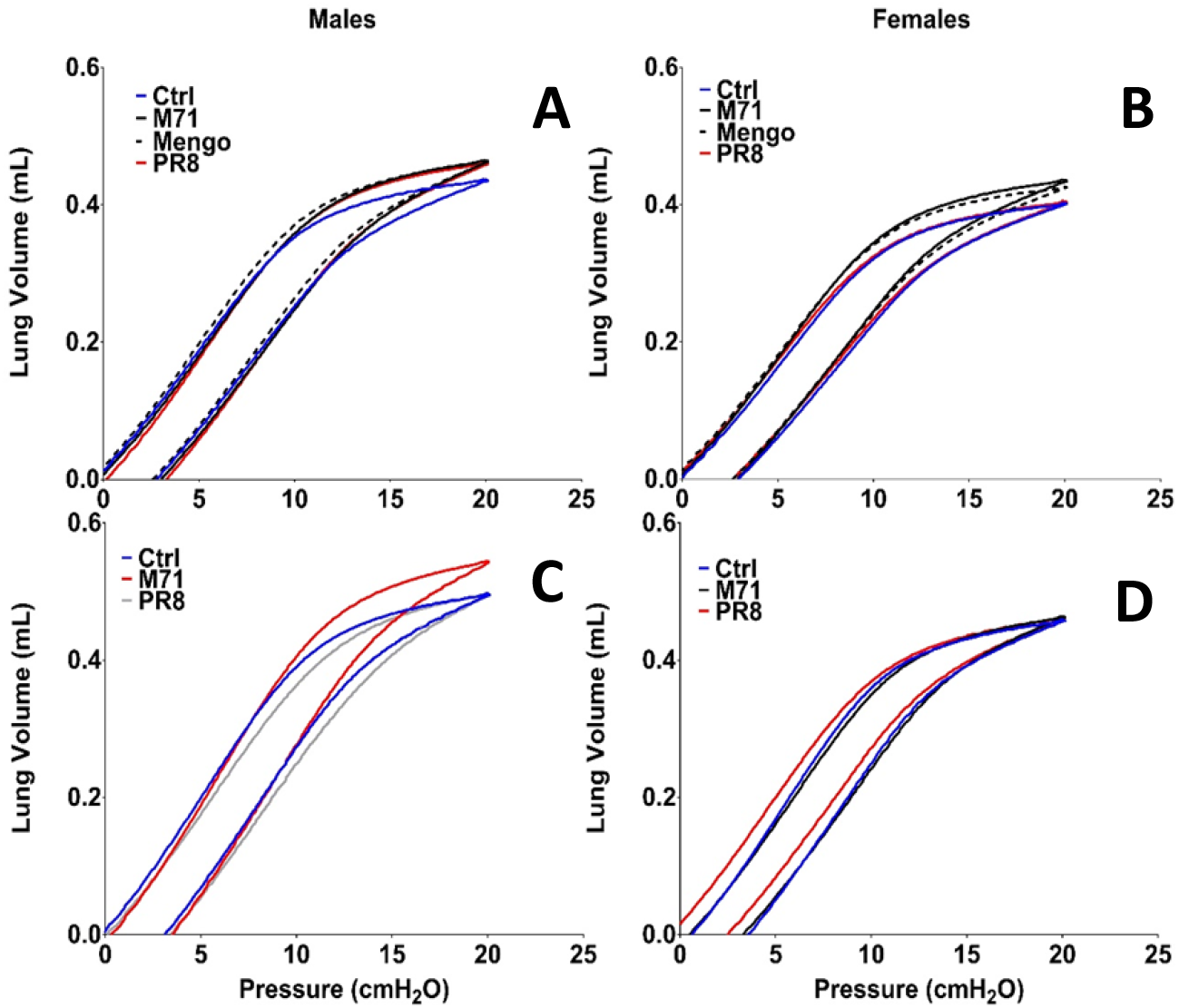
The average third deep ventilation of each viral infection and sex is shown. **A)** and **B)** refer to male and female mice from the first experiment, respectively. **C)** and **D)** refer male and female mice from the follow-up experiment, respectively.

PV curves were also utilised to extract compliance, specific compliance, Vmax or K (see methods) from which more information regarding the disease model can be ascertained. We found that infection had no effect on compliance (p = 0.775), specific compliance (p = 0.290), Vmax (p = 0.279) or K (p = 0.308) for male or female mice (Table 4). However, there was an effect of sex on compliance (higher in males, p = 0.004) and Vmax (higher in males, p < 0.001) but not specific compliance (p = 0.594) or K (p = 0.116). There was no significant interaction between early-life viral infection and sex on compliance (p = 0.441), specific compliance (p = 0.796), Vmax (p = 0.714) or K (p = 0.896).

**Table 4:** Compliance, specific compliance, Vmax, and K were calculated from the individual PV loops from the initial study. P < 0.05 was considered significant and this was denoted by an asterisk (*).

| Male |  |  |  |  |
| --- | --- | --- | --- | --- |
| Measurement | Ctrl (n = 5) | M71 (n = 14) | Mengo (n = 11) | PR8 (n = 15) |
| Compliance (mL/cmH<sub>2</sub>O)* (*p* = 0.004) | 0.04 ± 0.00 | 0.04 ± 0.00 | 0.04 ± 0.00 | 0.04 ± 0.00 |
| Specific Compliance (cmH<sub>2</sub>O)<sup>-1</sup> | 0.40 ± 0.18 | 0.36 ± 0.07 | 0.33 ± 0.11 | 0.44 ± 0.23 |
| Vmax (mL)* (*p* < 0.001) | 0.44 ± 0.05 | 0.46 ± 0.04 | 0.47 ± 0.03 | 0.46 ± 0.05 |
| K (1/cmH<sub>2</sub>O) | 0.80 ± 0.05 | 0.78 ± 0.03 | 0.81 ± 0.03 | 0.79 ± 0.05 |
| Female |  |  |  |  |
| Measurement | Ctrl (n = 9) | M71 (n = 7) | Mengo (n = 5) | PR8 (n = 5) |
| Compliance (mL/cmH<sub>2</sub>O) | 0.03 ± 0.00 | 0.04 ± 0.00 | 0.04 ± 0.00 | 0.03 ± 0.00 |
| Specific Compliance (cmH<sub>2</sub>O)<sup>-1</sup> | 0.39 ± 0.15 | 0.36 ± 0.06 | 0.33 ± 0.10 | 0.35 ± 0.14 |
| Vmax (mL) | 0.40 ± 0.04 | 0.44 ± 0.03 | 0.43 ± 0.02 | 0.40 ± 0.05 |
| K(1/cmH<sub>2</sub>O) | 0.80 ± 0.03 | 0.80 ± 0.03 | 0.81 ± 0.04 | 0.81 ± 0.04 |

In the follow-up study, early-life viral infections had no effect on compliance (p = 0.056), specific compliance (p = 0.387), Vmax (p = 0.085) or K (p = 0.172) for both male and female mice (Table 5). There was no effect of sex on compliance (p = 0.051), specific compliance (p = 0.723) or K (p = 0.293). However, sex was associated with Vmax (higher in males p < 0.001). There was no significant interaction between early-life viral infection and sex on compliance (p = 0.069), specific compliance (p = 0.338), Vmax (p = 0.096) and K (p = 0.231).

**Table 5:** The compliance, specific compliance, Vmax, and K for male and female mice extracted from the PV curves in the follow-up study. P < 0.05 was considered significant and this was denoted by an asterisk (*).

| Male |  |  |  |
| --- | --- | --- | --- |
| Measurement | Ctrl (n = 8) | M71 (n = 15) | PR8 (n = 8) |
| Compliance (mL/cmH<sub>2</sub>O) | 0.04 ± 0.00 | 0.05 ± 0.00 | 0.04 ± 0.01 |
| Specific Compliance (cmH<sub>2</sub>O)<sup>-1</sup> | 0.40 ± 0.15 | 0.69 ± 0.72 | 0.55 ± 0.29 |
| Vmax (mL)* ($p < 0.001$) | 0.50 ± 0.03 | 0.55 ± 0.04 | 0.49 ± 0.09 |
| K (1/cmH<sub>2</sub>O) | 0.79 ± 0.05 | 0.75 ± 0.05 | 0.73 ± 0.09 |
| Female |  |  |  |
| Measurement | Ctrl (n = 6) | M71 (n = 6) | PR8 (n = 6) |
| Compliance (mL/cmH<sub>2</sub>O) | 0.04 ± 0.00 | 0.04 ± 0.00 | 0.04 ± 0.00 |
| Specific Compliance (cmH<sub>2</sub>O)<sup>-1</sup> | 0.50 ± 0.18 | 0.66 ± 0.39 | 0.41 ± 0.24 |
| Vmax (mL) | 0.46 ± 0.04 | 0.45 ± 0.05 | 0.47 ± 0.03 |
| K (1/cmH<sub>2</sub>O) | 0.79 ± 0.04 | 0.75 ± 0.05 | 0.79 ± 0.06 |

### Lung function at functional residual capacity (FRC)

FOT partitions the respiratory system into multiple compartments and therefore has superior precision to the snapshot perturbation technique. We used FOT to measure R_aw_, G, H and ɳ at functional residual capacity (FRC). There was no significant effect of infection (M71, PR8, Mengo) on any parameter of lung function at FRC for either male or female mice (p > 0.05; Table 6).

**Table 6:**
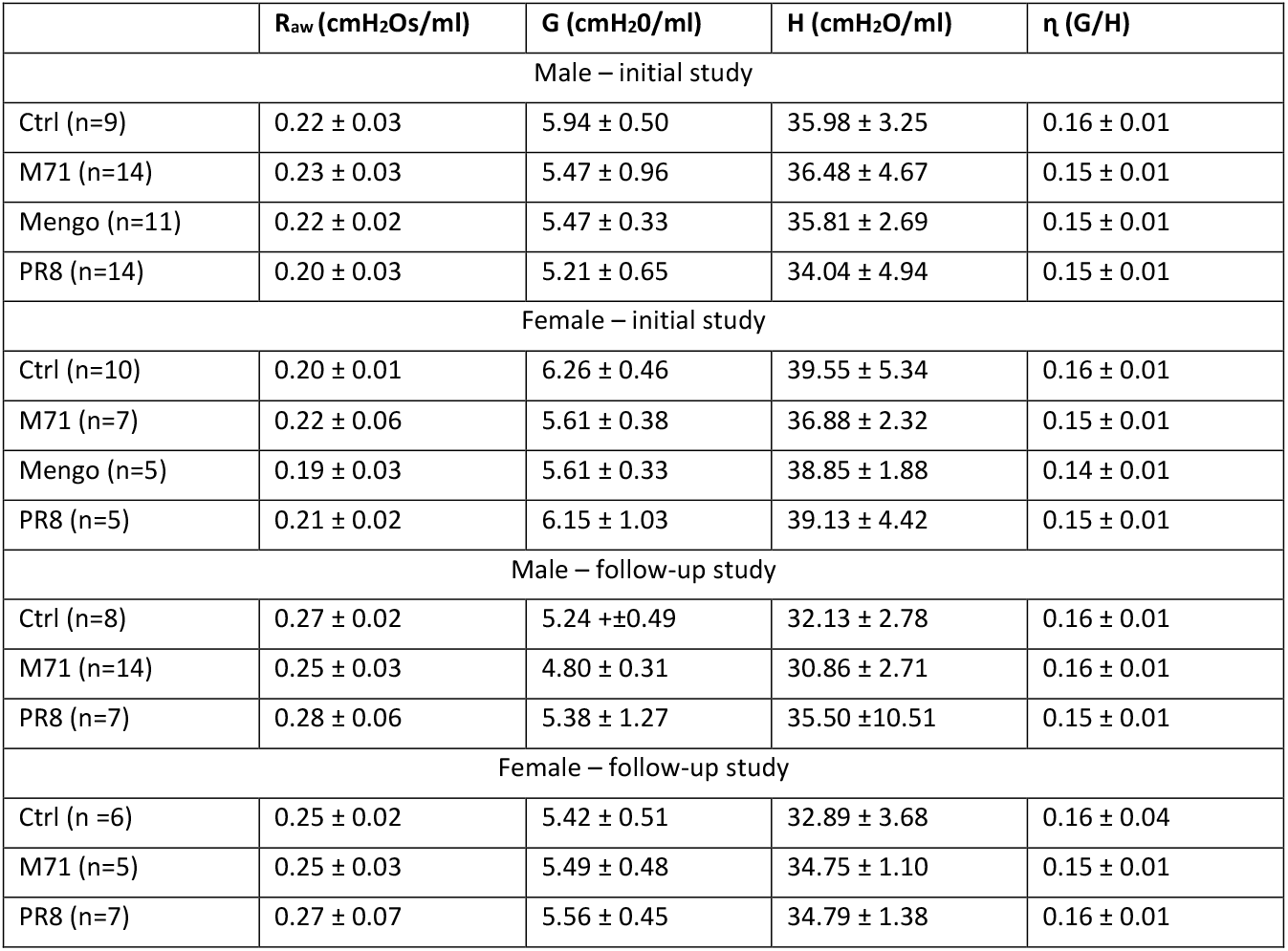
Lung function data for the R_aw_, G, H and η at FRC for each experimental group, in both sexes and experiments. Data presented are mean ± SD.

### Responsiveness to methacholine

MCh challenge was employed to measure airway hyperresponsiveness (29). We found that male and female mice infected with M71 or PR8 were more responsive to MCh, when compared to control, for R_aw_ (Figure 3; p < 0.001 for all comparisons). Moreover, M71 and PR8 infection also had an effect on G (males p < 0.001, females p < 0.001) and ɳ (males p <0.001, females p = 0.006) in both sexes. Furthermore, PR8 but not M71 had significant effects on H for both sexes (males p <0.001, females p <0.001, Figure 3). In contrast, male mice infected with Mengo had increased R_aw_ (p < 0.001), whereas female mice did not relative to their respective controls (p = 0.220). Moreover, there were also differences in G (p <0.001), H (p = 0.001), and ɳ (p < 0.001) in male but not female mice infected with Mengo. There was no effect of sex (p > 0.087) on R_aw_, G, H and ɳ, nor an interaction between sex and early-life viral infections (p > 0.551 for all lung function parameters). MCh sensitivity for all lung function parameters measured was not affected by infection (p = 0.074) or sex (p = 0.156) (Figure 5A and 5B).

**Figure 3:**
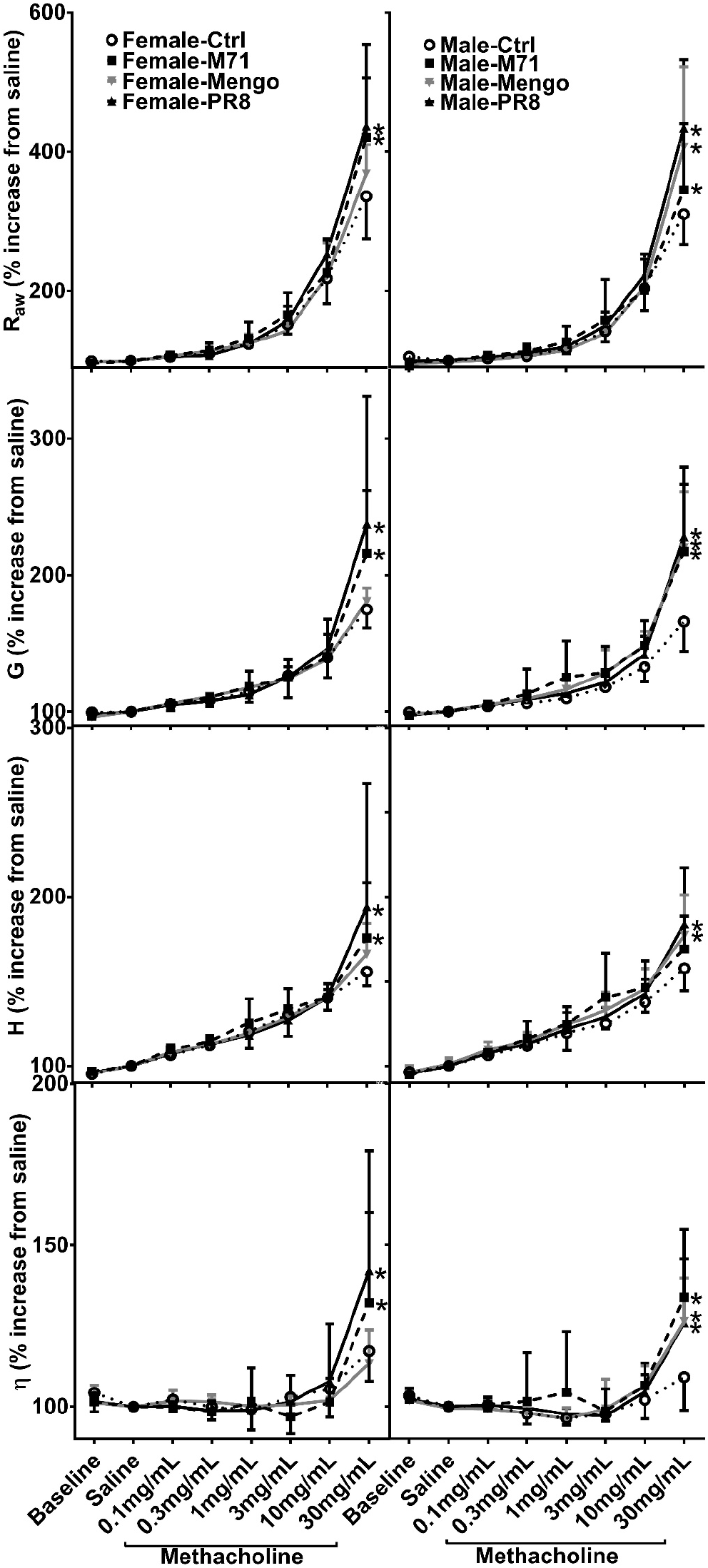
Dose response curves for female (left panels) and males (right panels). In every plot, Ctrl (female: n = 10; male: n = 9) is represented by a circle and a dashed line, M71 (female: n = 7; male: n = 14) by a square and a dashed line, Mengo (female: n = 5; male: n = 11) by a grey triangle and a dashed line and PR8 (female: n = 5; male: n = 14) by an inverted triangle and a solid line. A p < 0.05 was considered significant (* denotes significance at 30mg/mL MCh dose). Data are mean ± SD.

Similar trends were observed in the follow-up experiment, where R_aw_ was significantly elevated in male and female mice infected with M71 and PR8, when compared to control (Figure 4, p < 0.001). Furthermore, PR8 but not M71 significantly elevated G (male p < 0.001, female p = 0.008) in response to MCh, in both sexes, when compared to control. Moreover, M71 (male p = 0.214, female p < 0.001) and PR8 (male p = 0.624, female p < 0.001) had significant effects on H in female but not male mice. In contrast, male but not female mice infected with M71 (male p < 0.001, female p = 0.977) or PR8 (male p < 0.001, female p = 0.103) were more responsive to MCh, for ɳ, when compared to respective controls. We also found that female PR8 mice required less MCh before a significant change was detected in R_aw_ (REC125 p = 0.024), while M71 only affected H (HEC125 p = 0.044) (Figure 5C and 5D). All other parameters were not impacted by infection and sexes (p > 0.05 for all).

**Figure 4:**
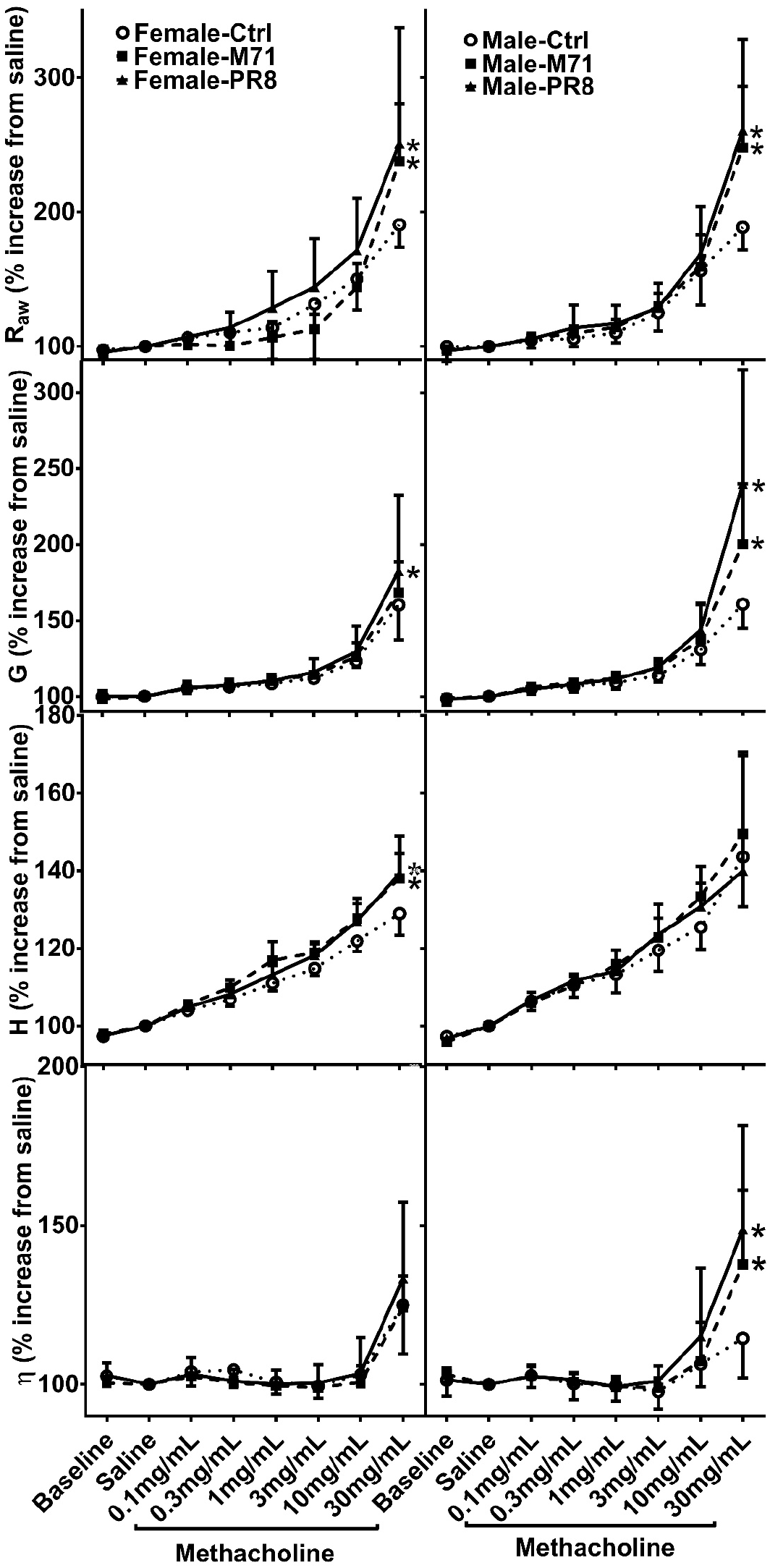
Dose response curves for female (left panels) and males (right panels). In every plot, Ctrl (female: n = 6; male: n = 8) is represented by a circle and a dashed line, M71 (female: n = 5; male: n = 13) by a square and a dashed line and PR8 (female: n = 7; male: n = 7) by a triangle and a solid line. A p < 0.05 was considered significant (* denotes significance at 30mg/mL MCh dose). Data are mean ± SD

**Figure 5:**
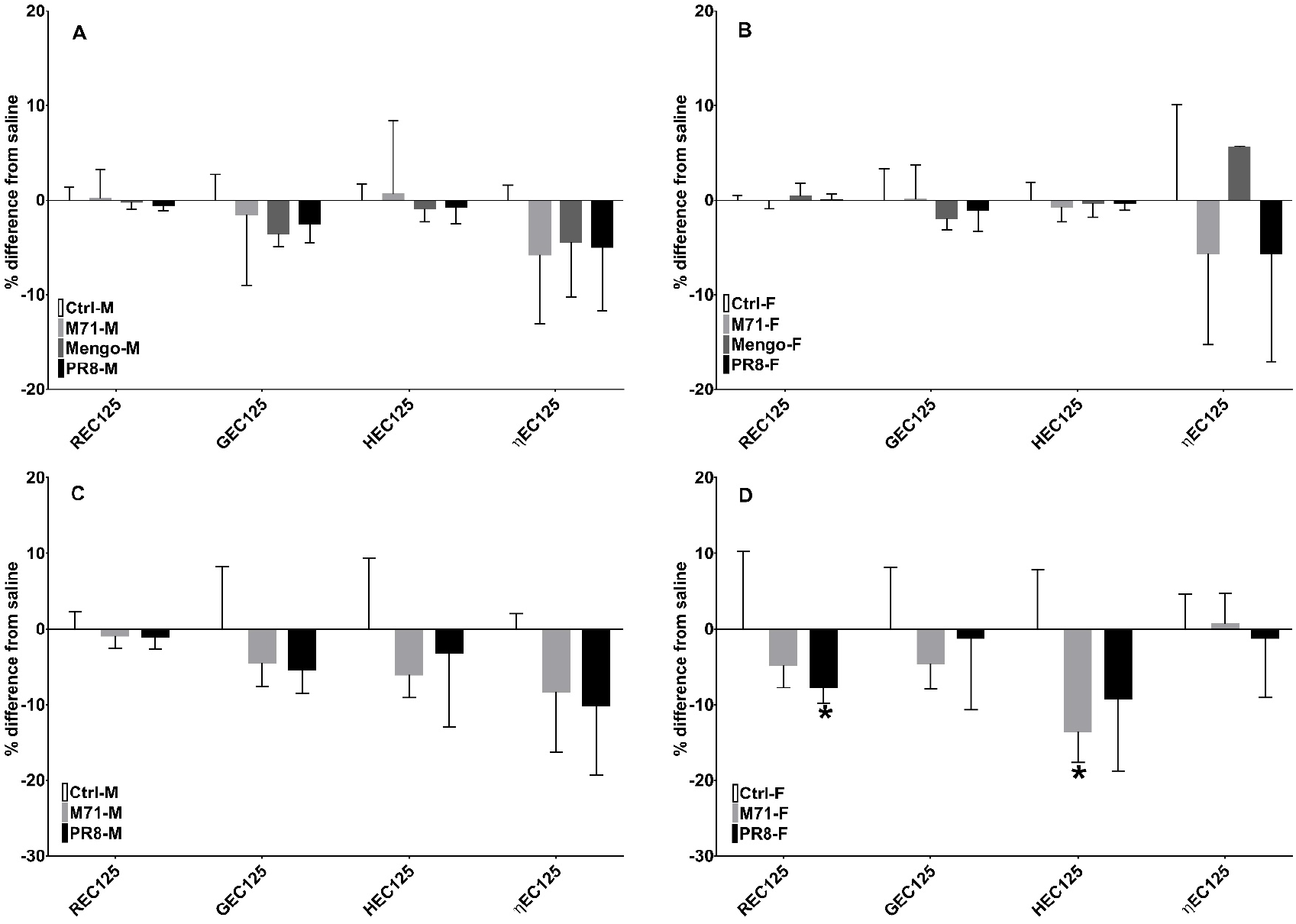
Methacholine (MCh) dose required to increase R_aw_ (REC125), G (GEC125), H (HEC125) and ɳ (ɳEC125) from saline by 25% for each experimental group. The infection groups were compared to control in both sexes and experiments. Males are grouped on the left and females grouped on the right. **A** and **B** panels refer to the MCh sensitivity in the initial experiment while **C** and **D** refer to MCh sensitivity in the follow up experiment. A p < 0.05 was considered significant and this was denoted by an asterisk (*).

### Comparison between initial study and follow-up study conducted in separate animal house facilities

We performed statistical comparisons on lung function at FRC between the initial study and the follow-up study to determine if the baselines differed between the experiments. The data showed there was no impact on R_aw_ (male: p = 0.262, female: p = 0.176), G (male: p = 0.249, female: p = 0.356), H (males p > 0.05, females p >0.05) or ɳ (male: p = 0.600, female: p = 0.553; Table 6). Moreover, early-life viral infection did not affect R_aw_ (male: p = 0.113, female: p = 0.286), G (male: p = 0.271, female: p = 0.498), H (males p > 0.05, females p > 0.05) or ɳ (male: p = 0.530, female: p = 0.233).

Then we performed a statistical analysis on MCh responsiveness at the highest dose (30mg/mL) on the lung function parameters (R_aw_, G, H and ɳ) between both locations. This would allow us to determine if there are differences in MCh responsiveness between experimental groups and sexes. We found that R_aw_ of Ctrl (males p = 0.029, females p = 0.031), M71 (males p = 0.012, females p = 0.006) and PR8 (males p < 0.001, females p = 0.007) mice in the initial study were more responsive to MCh than their counterparts in the follow up study. In contrast, tissue damping of female mice was significantly higher in the initial study than the follow up study (p = 0.037). While tissue elastance of male PR8 mice (and no other experimental group or sex) was significantly higher in the initial study than the follow-up study (p = 0.002). Moreover, hystersivity was not affected by the change in the animal facilities (males p = 0.060, female p = 0.988).

## Discussion

Early life represents a crucial period of heightened plasticity where deleterious environment exposures exemplified by respiratory viral infections can alter developmental and functional trajectories, leading to chronic diseases that persist throughout adulthood. Here, we tested the hypothesis that a single exposure to a viral pathogen on day seven of life (equivalent to first year in humans) could perturb the normal development of airway tissue leading to subsequent deficits in adult lung function. A key finding was that multiple viruses (M71, PR8, Mengo) could induce an AHR phenotype that was present in adulthood. Of particular interest, variations in these findings were observed with respect to virus type and sex. Notably, early-life Influenza M71 or PR8 induced AHR in both male and female mice, whereas induction of AHR following early-life Mengo was restricted to males. Cellular inflammation in BAL following M71 infection was characterised by peak inflammation at three days post-infection that waned soon after, as reported previously (29). Conversely, increased cellular inflammation post Influenza PR8 and Mengo infection was not observed in our study, when assessed 3-, 5- and 14-days post infection. Furthermore, the viral infections did not affect the somatic growth of male and female mice. In summary, we have demonstrated that a single respiratory viral infection on day seven of life leads to AHR in adulthood. These findings support the paradigm of the early origins of AHR, which has been reported to be present in almost one in five individuals (34). Furthermore, this model can be utilised for mechanistic studies to further our understanding of how viral infections perturb lung development and function over the life course.

Longitudinal studies in humans have demonstrated that respiratory viral infections in early life are associated with the development of chronic respiratory diseases such as asthma and chronic obstructive pulmonary disease (COPD) in later life (35). Moreover, in susceptible individuals, recurrent infections are associated with deficits in lung growth (children) and accelerated lung function decline (adults) (36). However, it is not possible to determine from observational studies in humans if viruses directly damage the airways driving disease development or alternatively unmask individuals with a pre-existing susceptibility that were already on a trajectory towards disease. Moreover, in human studies, it is difficult to determine the relative importance of the very first respiratory viral infection, which may occur during the neonatal period when the immune and respiratory systems are maximally susceptible, versus subsequent infections at later developmental stages. Notably, our finding that a single exposure to a viral pathogen on day seven of life leads to AHR in adulthood sheds light on the early origins of this phenotype. For instance, Collins et al. (34) investigated the risk factors associated with bronchial hyperresponsiveness (BHR) in teenagers and found BHR to be very common affecting almost 1 in 5 individuals. Moreover, the majority of individuals with BHR did not have current asthma or wheeze. It was also demonstrated that BHR was much more common in girls than boys. Interestingly, this finding could not be explained by the known risk factors of lower lung function (as girls tend to have larger airways than boys) or atopic status (with atopy being more common in boys), suggesting the hypothesis that viral infections may play a major role in explaining a significant proportion of the phenotypic variance. We assessed baseline lung function using the snapshot perturbation technique and FOT, with the hypothesis that viral infections perturb lung function at functional residual capacity. We did not observe any differences in the parameters measured using either the snapshot technique (R, E, C) or FOT (R_aw_, G, H, ɳ). In contrast, when the mice were challenged with increasing doses of MCh, we observed marked differences between the infected and control groups in all lung function parameters. Therefore, our data suggest that respiratory viral infection on day seven of life can induce AHR in adulthood in the absence of changes to baseline lung function. We also found that virus type had distinct effects on parenchymal lung mechanics (G and H). Influenza PR8 and M71 infection led to hyperresponsiveness to MCh in terms of tissue damping and tissue elastance irrespective of sex, while Mengo was found to only affect these parameters in male mice. Viral infection also affected lung homogeneity, indicated by the increase in hystersivity at the highest MCh dose. This suggests that peripheral lung resistance is increasing disproportionately to lung elastance as a consequence of early life infection. Tissue damping is a measure of the dissipative properties of the lung (37) while tissue elastance is a measure of elastic recoil (38). Both mechanical parameters are considered when evaluating lung health growth. For instance, healthy lung development in infants is marked by low air resistance in the central and peripheral airways of the lung in humans (39) and mice (40), whereas our data show these parameters are altered as a result of the infection, leading to increased air resistance in the central and peripheral airways.

RSV, RV, and Influenza are the three most common pathogens causing respiratory infections in children (41). As these viruses are readily encountered in nature, children are bound to experience multiple infections during development, and therefore it is difficult to determine the impact of the very first infection relative to subsequent infections on lung development. Our data demonstrates that a single infection on day seven of life was sufficient to induce AHR in adulthood. Moreover, this phenotype was induced by both PR8 and M71, even though the former virus is much more virulent than the latter (23). Together, this suggests that age of first infection may be more important than virulence (41, 42).

Males and females are known to experience variations in disease susceptibility and severity to viral infections (43). Accordingly, the interactions between sex hormones and the developing immune system and lung are important determinants of disease. For instance, in our study, we found that male mice infected with Mengo developed AHR phenotype in adulthood while female mice did not. These data are consistent with observations in human studies that have demonstrated that young boys are at a greater risk of rhinovirus-induced asthma exacerbations compared to girls (44).

Weight loss and inflammation are cardinal features of viral infections in mice. We found that viral infections did not lead to significant weight differences between experimental groups and sexes. This suggests that the viral infections have not hindered somatic growth and therefore perturbations observed in lung function parameters (R_aw_, G, H and ɳ) may be a direct consequence of the viral infections. Sex differences were noted in delta weights, with males weighing more than females regardless of experimental group, and similar findings have been published in previous studies (45, 46). Although cellular inflammation is an expected consequence of viral infections (47) (often characterised by an increase in total cell counts) we only observed cellular inflammation in BAL following M71 infection but not the other viruses. Our interpretation of these data is that the time points we sampled did not capture peak inflammation for Mengo (48) or PR8 (49) in mice at seven days of age, although further studies are required to address this issue.

Our study has limitations that should be acknowledged. Firstly, we focused on M71, PR8, and Mengo, but we acknowledge that RSV is an important pathogen encountered in early-life and was not included in our study. Secondly, we focused on BALB/c mice as they are Th2-skewed and are more susceptible to developing asthma-like traits. Including other strains for comparison would be important for understating the role of genetic background in disease susceptibility. Thirdly, were performed a single infection on day seven of life. Future studies could investigate the role of age of first infection on disease development and/or the impact of multiple infections. Notwithstanding these limitations, we have established an experimental mouse model of early-life respiratory viral infection that results in lung function impartment in adults. Our findings provide new insight into the early origins of AHR and the model provides a tractable experimental system for immunological and molecular studies to dissect the underlying mechanisms.

## Acknowledgments

This study was supported by the NHMRC project grant 1129996 and the Research Training Program Scholarship. We would like to extend our thanks to the staff at the Telethon Kids Institute Bioresources Centre for their assistance in this study.

